# Molecular Characterization of *Tephromela atra* and *Tephromela grumosa* from Bozdag Mountain Based on ITS1 rDNA Sequences Analysis

**DOI:** 10.1101/2024.01.04.574211

**Authors:** Awad Alzebair, Meral Yilmaz Cankiliç, Mehmet Candan, İlker Avan

## Abstract

The lichens *Tephromela atra* and *Tephromela grumosa* were studied in detail using various techniques, including morphological identification, biochemical taxonomy (chemotaxonomy), anatomical, and molecular characterization. The molecular characterization of *Tephromela atra* and *Tephromela grumosa* (Pers.) Hafelner & Cl. Roux was analyzed using internal transcribed spacer rDNA (ITS1) genes and the sequences obtained were compared with the ITS sequences of the other *Tephromela* species in GenBank (NCBI). The sequences were analyzed against reference sequences from the GenBank databases using the programs; BLAST, ClustalW, and MEGA11. The phylogenetic relationship was evaluated. The results show that some samples of *Tephromela atra* obtained in this study are closer to the same species isolated from Greece, while other samples are closer to other samples isolated from Italy and Austria. Moreover, the *Tephromela grumosa* isolated from Eskisehir Bozdag Mountain form a separate branch closer to the *Tephromela grumosa* samples isolated from Italy. Our study shows that lichens are an important source of biodiversity in Türkiye, which has a high species richness in Anatolia.

## INTRODUCTION

Lichens are a symbiotic partnership between green algae or cyanobacteria and fungi, which join together to form a single thallus, called lichen[1]. The fungal partner of the lichen is called the mycobiont, and the photosynthetic partner (cyanobacteria or algae) is called the photobiont. The mycobiont is responsible for providing water, carbon dioxide, minerals, and protection for the lichen thallus[2]. The photobiont supplies organic compounds and oxygen produced by chlorophyll to the structure of the lichen thallus[3]. Since the fungal partner plays the main role in the life of a lichen symbiosis, lichens are sometimes referred to as “lichenized fungi”. According to some researchers, lichens are therefore an example of the controlled parasitism that fungi establish on algae [4, 5].

Most lichen species were defined in the 1800s and the first half of the last century by the concept of phenotypic species based on monomorphism [6]. The differences in morphological characters and, to a certain extent, anatomical features were used to differentiate between species. These differences are environmental rather than genetic. These taxonomic studies have led to an accumulation of many synonymous species. As a result, biographic origin is of great importance and most allopathic populations are named as separate species. Conventional diagnostic keys for lichen diagnostics work according to the morphological, anatomical, and more rarely, chemical characteristics of the organisms. Originally, the differences in morphological and anatomical characteristics between different species may not have been sufficient for accurate identification of the organisms. In addition, organisms living in different environments may differ greatly from each other, so that organisms of the same species show different morphological and anatomical characteristics. Therefore, the genotype is more stable than the phenotype. Morphologically emerged diversity constitutes only a small percentage of genetic diversity and a large ratio of genetic diversity [7-10]. Some of the morphological characteristics are not reflected. Although the information obtained from morphological characteristics is used to identify genetic differences. The interaction between genotype and environment is not always sufficient because a character is identified by more than one locus and a gene affects more than one character[11]. In lichens, the concept of chemical species is also used to identify numerous taxa that cannot be distinguished morphologically[12]. However, this situation was first criticized by authors who thought that it was necessary to define the morphological characters[13, 14]. Systematic studies of lichens, as with other groups, are classically based on characterization by morphological characters. In addition to morphological characters, chemotaxonomic methods are still in use [6]. However, the fact that these identification studies do not fully define the lichen or plant is a problem. For this reason, systematic categories in recent years have led taxonomic studies to focus more on molecular methods, and molecular techniques and gene markers are the answer. Molecular markers have come to the fore in these molecular studies[7-9, 12].

Phylogenetic species concepts based on molecular characters are used to identify species within species complexes and to show how phenotypic characters in later generations develop slowly in lichenized fungi[15]. When studying genetically isolated species over a sufficiently long period, even a single locus can distinguish species in terms of fixed phenotypic characters. However, molecular characters such as phenotypic characters may be polymorphic within phylogenetic species and between sister species of species complexes. When a single locus cannot resolve species associated with phenotypic characters or biography, multiple loci should be used to accurately identify species. Most recent studies are based on nuclear ribosomal DNA (nrDNA) sequence information[16]. Several lichen species have been investigated in molecular phylogenetic studies[17-24]. However, there are only a few studies on *Tephromela atra*, and most of these studies were carried out at the University of Trieste in Italy by Lucia Muggia, for example, an early study by Muggia et al., 2008[25], reported extremotolerant fungi from alpine rock lichens, which include *Tephromela atra*, and their phylogenetic relationships[25-31].

As the literature search revealed, there is no previous study showing the diversity and molecular characterization of *Tephromela M. Choisy* species in Eskisehir Bozdag Mountain. Thereby, this study aimed to molecularly identify two different species of the lichen genus *Tephromela M. Choisy* isolated from Bozdag Mountain in Eskisehir and to determine their phylogenetic relationship to other *Tephromela* lichens. The results of the present study indicate that some of the *Tephromela atra* samples are more similar to species that were previously isolated from Greece and others show similarity to the ones that were isolated from Italy and Austria. In addition, *Tephromela grumosa* isolated from Eskisehir Bozdag Mountain forms a separate branch that is closer to the Italian *Tephromela grumosa* samples.

### Tephromela M. Choisy (1929)

*Tephromela M. Choisy* the name comes from the Greek word “tephra”, meaning ash, and “mela”, meaning black, and thus refers to the black color of the lichen thallus. The *Tephromela* species share diagnostic anatomical characters with the type species, *Tephromela atra* (Huds.) Hafellner & Kalb[32]. *Tephromela atra* and *Tephromela grumosa* examined in this study are shown in Figure 1.

**Figure 1.**
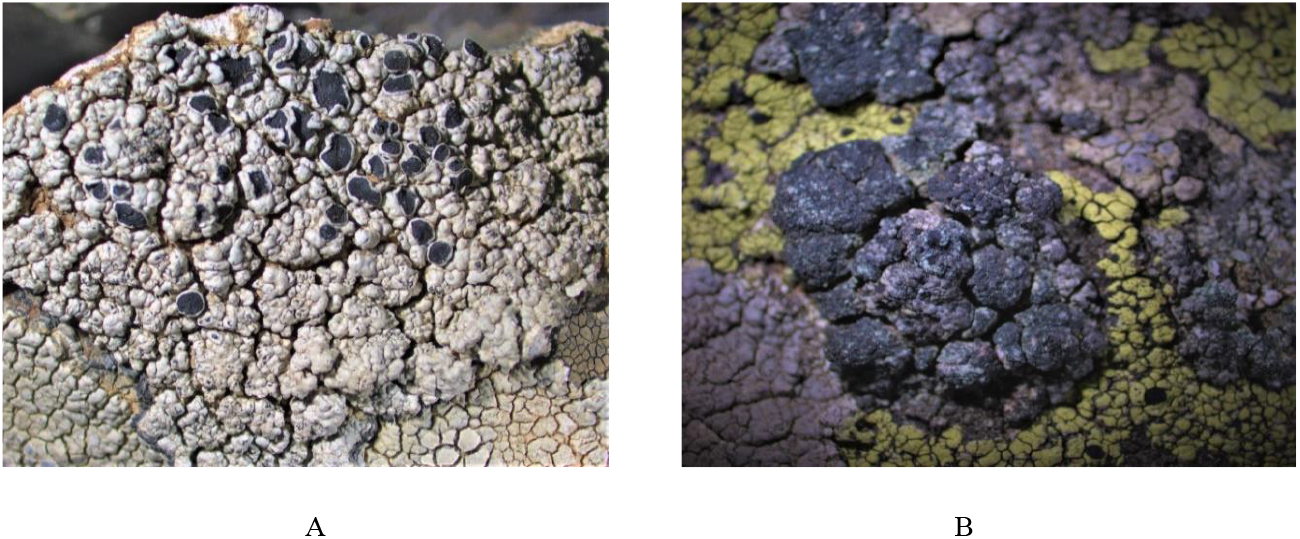
Microscopic images of *Tephromela* lichens collected from Eskisehir Bozdag Mountain (A: *Tephromela atra*; B: *Tephromela grumosa*).

### Distribution of *Tephromela* and study area

The genus *Tephromela* comprises around 25-30 species. One of the most common species is *Tephromela atra*. Various identification keys, flora books, and monographs were used to identify the species[1, 33-38]. *Tephromela atra* has a worldwide distribution and has been reported from Chile, Antarctica, Argentina, New Zealand, and Australia. It is unknown on the other side of the world. However, numerous other species have been reported, especially from Antarctica and Australia. In southern South America, the only other species reported besides *T. atra* is *T. austrolitoralis* (Zahlbr.), reported by Elix and Kalb, (2008)[39]. It resembles *T. atra* but has an interspersed hymenium. *Tephromela atra* has been recorded in most provinces of Türkiye in the last thirty years. It is reported for examples from Bursa, Sivas, Adana, and Ankara. The first studies that recorded *Tephromela atra* lichen from Bozdag Mountains in Eskisehir province belong to Özdemir, (1991)[40], Özdemir Türk, (2002)[41], John V, (2007)[42], Singer, et al, (2014)[43] and John and Türk, (2017)[44]. Bozdag Mountain is located north of the center of Eskisehir and forms the western end of the Sundiken Mountains. The southern slopes of the mountain start with an altitude of about 850 meters in the grasslands of Eskisehir, the highest point is 1534 meters, and the northern slopes descend steeply into the Sakarya Valley, about 200-250 meters.

## MATERIALS AND METHODS

### Collection of lichen samples

The lichens *Tephromela atra* and *Tephromela grumosa* (Figure 1) were collected from an open area on the siliceous rock in the Bozdag Mountain in Eskisehir province. Verification and morphological characteristics of *Tephromela atra* and *Tephromela grumosa* lichens were carried out using standard methods[35]. In order to facilitate the identification, lichen samples were collected from each locality together with the substrate on which they live. In this way, the type of *Tephromela* spp. on which the lichens developed could be determined in a laboratory study. To facilitate the identification of the lichen species, thallus pieces with structures responsible for the reproduction and dispersal of the lichens were preferably taken. The samples collected in the field study were wrapped with paper napkins and placed in paper bags. The location, date of collection, information about the substrate type, height, and coordinates measured with the GPS device were noted. Moist ones from the samples collected in the study field were dried in the laboratory at room temperature. To protect the identified samples from the negative effects of pests, they were kept in good condition. The samples were then placed in special herbarium envelopes of 12 × 17 cm size. Information about the lichen sample was noted on the label of the envelope. Finally, the samples are stored in the herbarium of Eskisehir Technical University, Department of Biology, Eskisehir, Türkiye.

### DNA Extraction and Amplification

DNA extraction was performed using a combination protocol between the NucleoSpin® Plant II kit from Macherey-Nagel and the DNeasy plant mini kit from QIAGEN. Approximately 2 g of each lichen sample was grinding and crushed with a mortar and pestle for distribution of the sample and convert the lichen samples into powder. Liquid nitrogen was then added to the tube and freeze the lichen samples for approximately 45 seconds and the samples were kept submerged in liquid nitrogen and disrupted for approximately 30 seconds at full speed and allowed the liquid nitrogen to evaporate. Then 20 mg of glass beads were added to each sample. The sample was mixed for 20 minutes using the vortex accessory from “Thomas Scientific”. All steps of DNA extraction are the same as in the kit manufacturer’s instructions with some differences: For cell lysis,400 μL of AP1 lysis buffer from the DNeasy plant mini kit plus 400 μL of PL1 buffer from the NucleoSpin® Plant II kit was used, the uses of these two buffers in the protocol is based on the established CTAB procedure. The general chemicals utilized in the lysis buffer are 2% CTAB, 0.04 M Tris, 0.04 M EDTA pH: 8, MgCl2, SDS, Triton X100, KCl, 0.8 M LiCl, NaCl, and other detergents. For DNA elution, 50 μL of the PE elution buffer from the NucleoSpin® Plant II kit was added and the tube was incubated at 65 °C for 5 minutes before centrifugation, repeating this step twice. The presence of DNA was determined by loading the DNA samples onto the gel using gel electrophoresis techniques. The gels were prepared from 1.5% agarose with 1x TAE buffer (Tris-acetate-EDTA). With RedSafe™ Nucleic Acid Staining Solution (20,000x) dye was added at a final concentration of 0.5 mg/ml for visualizing the DNA fragments, Finally, the 1kb DNA ladder was loaded as a positive control, and samples were loaded on a gel for 70 minutes at 100 volts then agarose gel images transferred to the UV imaging system, whether banding and DNA sizes have been identified.

The phylogenetic study was analyzed with sequence data of the ITS1 gene region (Figure 2) and the ITS1 was amplified using PCR technique with the primers ITS1 (CTTGGTCATTTAGAGGAAGTA), and ITS4 (TCCTCCGCTTATTGATATGC).

**Figure 2.**
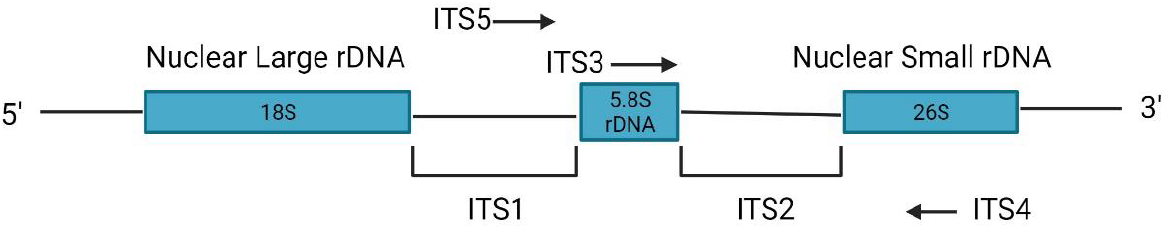
Diagrammatic representation of the Internal Transcribed Spacer (ITS) regions of 18S-26S nuclear ribosomal DNA (nrDNA). The positions of the ITS genes group used in this study for amplification and sequencing are indicated by an arrow.

**Table 1.**
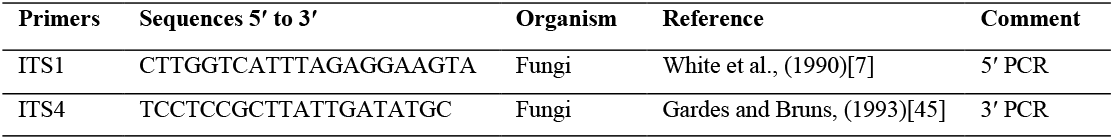
PCR ITS primers sequences.

Amplification of the ITS region in the genomic DNA of the lichen was performed with the primer containing the corresponding region under appropriate PCR conditions. The PCR reaction mixture used; Within 25 μL volume: 0.5 μL ∼ 200 ng DNA, 2.5 μL 10X reaction buffer, 1 μL 0.1 mM dNTPs, 0.75 μL 10μM ITS1 primer, 0.75 μL 10μM ITS4 primer, 0.5 μL Taq DNA polymerase and will complete the total volume to 25 μL PCR reaction was performed by adding 17 μL sterile distilled water. PCR reactions are initiated on the Progene-Techne PCR instrument under the following conditions: 94 ºC 5 min. pre-denaturation, 94 ºC 45 sec., 56 ºC 30 sec., 72 ºC 2 min. was performed as the final elongation phase. The DNAs amplified under these PCR conditions were run on agarose gel (1.5 % agarose gel, 100 volts, 30 minutes) and visualized by gel electrophoresis. Banding and negatives were checked after UV imaging. Since it was observed that the PCR products in the gel images could not be obtained in the desired purity in the gel images obtained by PCR, the reactions were repeated several times under different PCR optimization conditions. The MgCl2 concentrations were changed and the DNA that showed the correct bands was stored at 4°C for purification.

### Sequencing, Alignment, and Phylogenetic Analyses

The target DNA region amplified by PCR was extracted from the gel and included in the sequence analysis. The sequences analyzed from a double chain were checked with the BLAST (Basic Local Alignment Search Tool) program using data from GenBank (National Center for Biotechnology Information) (www.ncbi.nlm.nih.gov). The percentage match of the sequences was determined, and the matching sequences were selected. The integrity of the data was confirmed by checking the compatibility of the bidirectional arrays within themselves and solving the possible read errors with the Cluster X program. After the sequences were sorted, the unstable regions were removed and evaluated as lost data when performing the analyses. Six samples for the ITS1 gene region were used for the molecular analyses. During this process, each base was carefully checked. The analyses were performed with the MEGA11 program. It was calculated by analyzing the ITS sequences. After obtaining the DNA sequences, the sequences were analyzed using reference sequences from the GenBank (NCBI) databases, using the BLAST, ClustalW, and MEGA11 programs. Multiple alignments of the downloaded sequences and the sequences obtained in our study belonging to the species were performed and determined by comparing them with the data in GenBank with the BLAST program. Based on the aligned sequences, a phylogenetic tree showing the grouping of species was generated using the MEGA11 program to understand the relationships between species and populations. For the phylogenetic tree of the ITS region, *Lecanora cenisia* (EU558540.1) and *Lecanora rupicola* (AY398707.1) were selected as the outer group.

## RESULTS

### DNA Extraction

Samples of *Tephromela* species were collected at various locations and identified using classical systematic methods. A total of six samples of *Tephromela atra* (Contig 2, 3, 5, 7) and *Tephromela grumosa* (Contig 4, 6) DNA were isolated. The names and codes of the species with DNA isolation are listed in Table 3. The isolation products were carried out in gel electrophoresis and the presence of DNA was determined from the band’s photo taken as a result of gel electrophoresis.

### Sequences and Molecular Analysis

As a result of the PCR, the samples of the target region were amplified, and the desired bands were purified by standard purification methods. The products obtained from the purification were sequenced by BM Labosis Company. The sequences of the species were aligned using the programs BioEdit, ClustalX 2.0, and MEGA6, and an analysis of the sequences was performed. Phylogenetic trees were generated using the maximum likelihood algorithm and the Nearest Neighbor Interchange (NNI) method in MEGA 11 (Figure 3). The distance matrix was calculated using the Jukes-Cantor correction and the validity of the tree topology was checked using the bootstrap method (1000 replicates). The accession numbers of the DNA sequences of *Tephromela atra* and *Tephromela grumosa* used in the creation of phylogenetic trees are given in Table 3. To compare the sequences of *Tephromela* species obtained in this study with other *Tephromela* species from other parts of the world that were sequenced and included in the database, the sequences were extracted and merged with other sequences downloaded from the GenBank database (www.ncbi.nim.nih.gov) as shown in Table 2. The alignment of the sequences was performed visually, as only a few gaps were present and easy to interpret. Insertion/deletion gaps were treated as missing data. According to the phylogenetic tree formed after the minimum evolution (ME) analysis (Figure 3), the species belonging to the *Tephromela* species were found to be divided into three main branches. *Lecanora cenisia* (EU558540.1) and *Lecanora rupicola* (AY398707.1) are the outgroup species in Minimum-Evolution (ME) analysis and has created a separate branch from all studied species. When comparing the differentiation of the studied *Tephromela atra* and *Tephromela grumosa* with the related genus, the species belonging to the same genus generally formed a branch against the main branch. According to the results of the molecular phylogenetic analysis in Figure 3, *Tephromela atra* (PP003331 and PP003332) was found to be closer to the same species isolated from Greece than the other samples obtained in this study, which are closer to other samples isolated from Italy and Austria. Furthermore, the phylogenetic tree shows that the sequences of *Tephromela grumosa* from Eskisehir (PP003333 and PP003335) form a separate branch closer to *Tephromela grumosa* samples isolated from Italy. In this study, the phylogenetic analysis of *Tephromela atra* and *Tephromela grumosa* isolated from Bozdag Mountain Eskisehir was evaluated for the first time in the literature, and it is expected that the results will be a source for future identification of lichen species by molecular phylogenetic methods.

**Table 2.** Origins, voucher code, and accession numbers of sequences that were downloaded from GenBank.

| Species | Voucher code | Accession No. | Origin |
| --- | --- | --- | --- |
| *Tephromela atra* | TSB 37121 | EU558649.1 | Greece |
| *Tephromela atra* | TSB 37456 | EU558659.1 | Greece |
| *Tephromela atra* | TSB 37930 | EU558690.1 | Greece |
| *Tephromela atra* | TSB 37922 | EU558686.1 | Greece |
| *Tephromela atra* | TSB 37919 | EU558685.1 | Greece |
| *Tephromela atra* | TSB 37942 | EU558614.1 | Greece |
| *Tephromela atra* | TSB 37940 | EU558609.1 | Greece |
| *Tephromela atra* | TSB 37924 | EU558688.1 | Greece |
| *Tephromela atra* | TSB 37914 | EU558682.1 | Greece |
| *Tephromela atra* | TSB 37910 | EU558680.1 | Greece |
| *Tephromela atra* | TSB 38695 | EU558616.1 | Italy |
| *Tephromela atra* | TSB 38698 | EU558619.1 | Italy |
| *Tephromela atra* | TSB 37936 | EU558605.1 | Italy |
| *Tephromela atra* | TSB 38697 | EU558618.1 | Italy |
| *Tephromela atra* | TSB 38699 | EU558620.1 | Italy |
| *Tephromela atra* | TSB 37928 | EU558689.1 | Italy |
| *Tephromela atra* | TSB 37137 | EU558655.1 | Austria |
| *Tephromela atra* | TSB 37903 | EU558676.1 | Austria |
| *Tephromela atra* | TSB 37917 | EU558684.1 | Greece |
| *Tephromela atra* | TSB 37923 | EU558687.1 | Greece |
| *Tephromela atra* | L1316 | KF730620.1 | Türkiye |
| *Tephromela atra* | L1320 | KF730621.1 | Türkiye |
| *Tephromela atra* | MB 0.004 | KX550110.1 | Türkiye |
| *Tephromela grumosa* | TSB 37070 | EU558628.1 | Italy |
| *Tephromela grumosa* | TSB 38686 | EU558625.1 | Italy |
| *Tephromela grumosa* | TSB 37082 | EU558641.1 | Italy |
| *Tephromela grumosa* | TSB 37881 | EU558673.1 | Italy |
| *Tephromela grumosa* | TSB 37079 | EU558635.1 | Italy |
| *Tephromela grumosa* | TSB 38687 | EU558626.1 | Italy |
| *Tephromela grumosa* | TSB 37080 | EU558636.1 | Italy |
| *Lecanora cenisia* | TSB 37464 | EU558540.1 | Italy |
| *Lecanora rupicola* |  | AY398707.1 | Austria |

**Table 3.** Names of the sample’s species, codes, origin, and accession numbers of DNA sequences used in the creation of phylogenetic tree.

| Species | Sequence ID | Accession No. | Location (Origin) |
| --- | --- | --- | --- |
| *Tephromela atra* | Contig-2 | PP003331 | Bozdag Mountain |
| *Tephromela atra* | Contig-3 | PP003332 | Bozdag Mountain |
| *Tephromela grumosa* | Contig-4 | PP003333 | Bozdag Mountain |
| *Tephromela atra* | Contig-5 | PP003334 | Bozdag Mountain |
| *Tephromela grumosa* | Contig-6 | PP003335 | Bozdag Mountain |
| *Tephromela atra* | Contig-7 | PP003336 | Bozdag Mountain |

**Figure 3.**
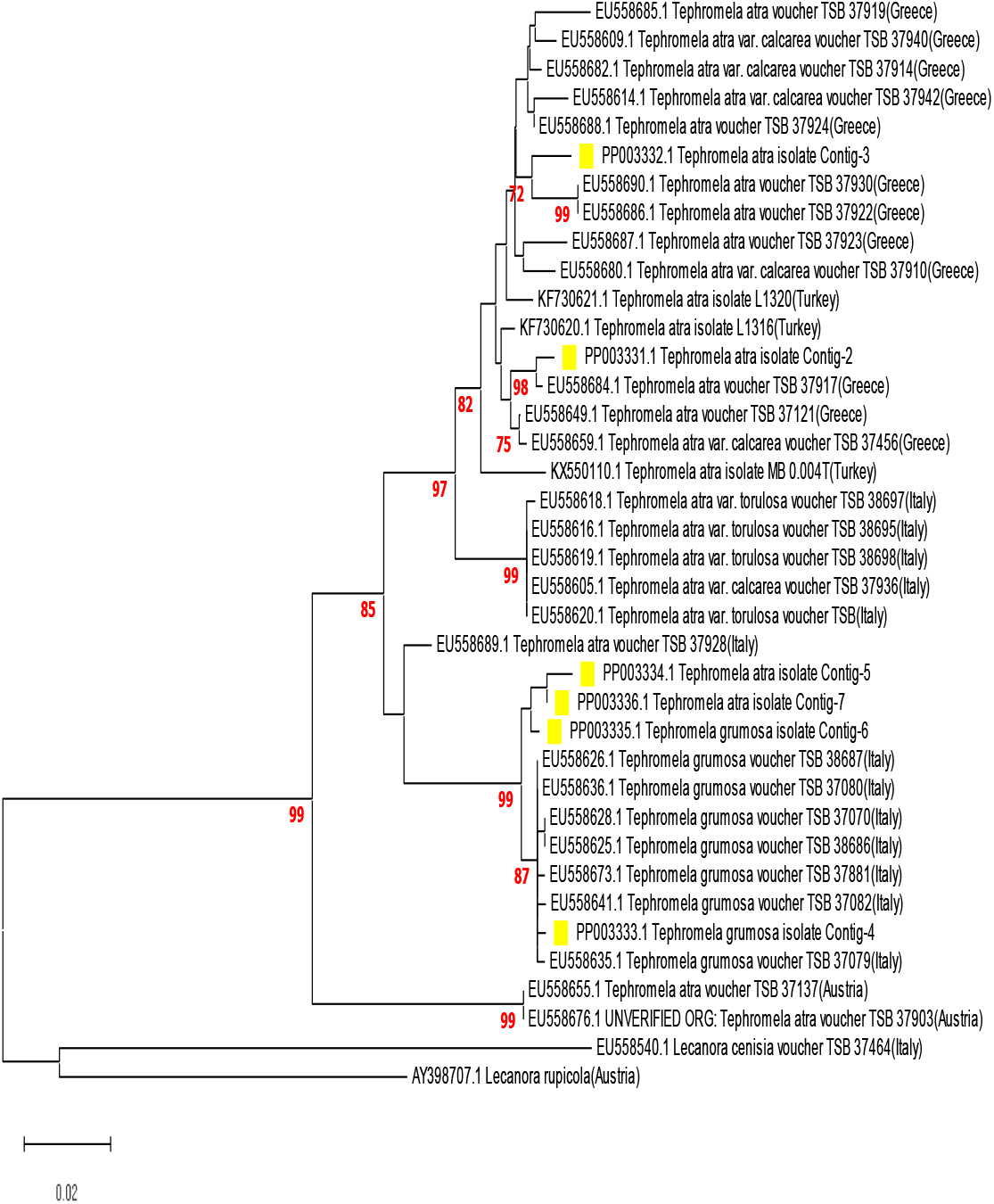
Molecular Phylogeny evolution analysis of *Tephromela atra* and *Tephromela grumosa* inferred from ITS region sequences data. The sample selected from GenBank is reported with their accession numbers. Newly obtained sequences in this study are in yellow. Scale bar represents the probable number of substitutions per site.

## CONCLUSION

To the best of our knowledge, the diversity and prevalence of *Tephromela* lichens in Türkiye have not been previously studied. This is the first report using elaborate techniques such as morphological taxonomy, morphological identification, biochemical taxonomy (chemotaxonomy), anatomical characterization, molecular identification, and phylogenetic techniques to study *Tephromela* species in Eskisehir. In Türkiye, many ITS sequence analysis information of different lichen species have been included in GenBank, but no phylogenetic studies on *Tephromela atra* and *Tephromela grumosa* species from Bozdag Mountain have been performed in the literature. In this study, the DNA sequence analysis and phylogenetic analysis methods were applied to determine the evolutionary relationships. In previous studies, various methods such as RAPD and AFLP were used for genotyping, but these techniques cause problems due to the symbiotic life of lichens. In this study, ITS sequences were collected from six samples of two different *Tephromela* species, *Tephromela atra* and *Tephromela grumosa*. The sequences were analyzed and compared with the data in GenBank. Methods based on PCR technology were used for genotyping because of its advantages. A highly informative phylogenetic tree was formed from the data obtained in this study. The ITS1 sequence analysis data showed the genetic similarities and differences in the lichens very well. The genetic distances observed in the phylogenetic tree indicate that the differentiation between the *Tephromela* species is quite high. The use of ITS1 sequence analysis, using only fungal-specific primers, has overcome these obstacles, and this method demonstrated in the study, is sufficient to reveal genetic differences in samples of the genus *Tephromela*. The results show that the genus *Tephromela* has a very informative phylogenetic tree consisting of many branches including *Tephromela atra* and *Tephromela grumosa* Species. Finally, this study shows that lichens constitute an important part of biodiversity in Türkiye which proves the richness of species in Anatolia. Furthermore, ITS diversity among species can help shed light on evolutionary variation among lichen species, especially *Tephromela* species.

## ABBREVIATIONS

*T. atra*: *Tephromela atra*
rDNA: Ribosomal DNA
ITS: Internal transcribed spacer region
ML: Maximum likelihood
ME: Minimum-Evolution
NNI: Nearest Neighbor Interchange
MSA: Multiple sequence alignment
CTAB: Cetyl trimethyl ammonium bromide
EDTA: Ethylenediaminetetraacetic acid
NCBI: National center for biotechnology information
PCR: Polymerase chain reaction
PE: Paired end
SD: Standard deviation

## ACKNOWLEDGEMENT

The authors thank Eskisehir Technical University, Scientific Research Projects Commission for funding the isolation of lichen acids and molecular studies (Project No. 19ADP089).

## Availability of Data and Materials

Assembled sequences are deposited in NCBI’s Sequence GenBank (accession numbers PP003331-PP003332-PP003333-PP003334-PP003335-PP003336).

## Authors’ Contributions

Conceptualization, A. A., and M. Y.; Methodology, A. A., and M. Y.; Validation, M. Y., I. A., and M. C.; Investigation, A. A., and M. Y.; Writing – Original Draft Preparation, A. A.; Writing – Review & Editing, A. A., M. Y., I. A. and M. C.; Supervision, M. Y.; Funding Acquisition, M. Y. All authors read and approved the final manuscript.

## Ethics Declarations

### Consent for publication

All authors confirmed the publication of the manuscript.

### Ethical approval

Our research does not need ethical approval.

### Human and animal rights

This article does not contain any studies on humans or animals subjects.

